# *Bacillus velezensis* SQR9 sways the rhizosphere community and its sociality toward cooperation and plant growth promotion

**DOI:** 10.1101/2022.10.11.511700

**Authors:** Yan Liu, Zhihui Xu, Weibing Xun, Polonca Stefanic, Tianjie Yang, Youzhi Miao, Nan Zhang, Ruifu Zhang, Qirong Shen, Ines Mandic-Mulec

**Affiliations:** Jiangsu Provincial Key Lab of Solid Organic Waste Utilization, Jiangsu Collaborative Innovation Center of Solid Organic Wastes, Educational Ministry Engineering Center of Resource-Saving Fertilizers, The Key Laboratory of Plant Immunity, Nanjing Agricultural University, 210095 Nanjing, Jiangsu, Peoples R China; Department of Microbiology, Biotechnical Faculty, University of Ljubljana, 1000 Ljubljana, Slovenia

## Abstract

According to the Lotka-Volterra competition model, intraspecific competition in plant-associated microbial communities should be stronger than interspecific competition. However, there is limited information on whether microbial communities follow this pattern and how disturbance by a newcomer affects them. Given the increasing popularity of probiotics, filling this knowledge gap could help guide future coexistence research. Here, we show that inoculation with a known probiotic, *B. velezensis* SQR9 shifts species co-occurrence patterns by decreasing the diversity of more distant species and promoting the growth of more closely related species, especially within the *Bacillus* community. By testing the sociality of *Bacillus* rhizosphere isolates, we then demonstrated that SQR9 increases the frequency of cooperative interactions in the *Bacillus* community, which may contribute to the promotion of plant growth. Finally, we provide an ecosystem framework comprising the strain’s genetic relatedness, metabolic niche space and social compatibility for the efficient and reliable assembly of *Bacillus* consortia. These findings shed new light on the ecological mechanisms underlying the beneficial effects of probiotics on host fitness.

## INTRODUCTION

The rhizosphere microbiome, also referred to as the second genome of the plant, can have a variety of beneficial effects on plant growth, nutrition and health^1, 2^. In the agricultural context, plant microbiomes have been proposed as the cornerstone of the next green revolution by improving crop performance while reducing undesired chemical remnants^3^. Moreover, microbial inoculants, commonly referred to as plant growth-promoting rhizobacteria (PGPR), have shown promise for improving plant growth, nutrition, and stress resistance^4,5^. In general, PGPR function by directly affecting the synthesis of phytohormones, enhancing nutrient uptake from soil and suppressing soil-borne plant pathogens^6, 7^. They also act indirectly by impacting the resident soil microbiome. To date, however, little research has been conducted on the effects of PGPR inoculants on the indigenous plant microbiota^8, 9^, and thus, the extent to which introduced PGPR inoculants affect the composition and function of the rhizosphere microbiome is largely unknown.

Rhizosphere competence and root colonization are two essential traits for the successful application of PGPRs as probiotics and biocontrol agents^10, 11^. For example, *Bacillus* and *Pseudomonas*, widely used as microbial inoculants, usually form robust biofilms on the plant root surface in the early stages after application^12^. Due to the intensive colonization of roots, these bacterial species can influence the composition of indigenous microbial communities by enriching microorganisms that are beneficial to host plant health^8, 13^, but empirical evidence is scarce. The application of PGPR is theorized to have the greatest negative impact on microbial species whose metabolic niches overlap^14, 15^. However, ecological interactions with other species may also alter the niche space of a given species^16^. According to theory and experiments, competition and antagonistic interactions reduce the niche space available to organisms^17^. In contrast, the theory predicts that mutualistic interactions expand the species’ niche space. However, empirical evidence is lacking on how the use of beneficial bacteria in the rhizosphere affects social trends in a community and its metabolic niche space^18^.

Social interactions are widespread in the microbial world, and cells can interact with each other in a variety of ways, resulting in either competition or cooperation between neighbours^19, 20^. PGPRs produce a variety of secondary metabolites, including bacteriocins^1, 21^, which can antagonize surrounding bacteria^20^, but less is known about their cooperative behaviors. For example, *Bacillus* spp., one of the soil organisms that exhibit complex social behaviors, such as cooperative swarming, preferentially merge their swarms with close relatives^22, 23^, a differential behavior toward relatives, known as kin discrimination. Specifically, within *B. subtilis*, kin soil isolates, defined as strains with at least 99.5% sequence identity at the housekeeping gene level, mostly merged their swarms and invaded a new territory as co-operators. In contrast, isolates with a nucleotide identity below 99.5% formed a visible boundary line at the site of encounter and did not enter a common swarm^24–26^. In the coincubation experiments, kin cells formed mixed biofilms on the plant root, whereas non-kin cells engaged in antagonistic interactions with one strain primarily colonizing the plant root surface^25^. Interestingly, associations of *B. subtilis* swarms with other species within the genus did not strictly follow this pattern, antagonizing closely related species and sometimes merging with more distantly related *Bacillus* spp^26^. Hence, relating phylogenetic shifts in the indigenous rhizo-*Bacillus* community to social interactions and community function remains a challenge, despite the importance of this information for a more efficient design of microbial inoculants.

Here, we fill this knowledge gap by using the well-known gram-positive PGPR strain *Bacillus velezensis* SQR9, a commercially available product (BIO™, China)^27^ with an outstanding ability to suppress soil-borne diseases and promote plant growth, especially cucumber growth^13, 27–32^. Through carefully designed pot experiments supported by bioinformatics of the bacterial community in the rhizosphere and in-depth functional analyses of the rhizosphere isolates, we show that *B. velezensis* SQR9 strongly affects the cooccurrence of the indigenous rhizosphere bacterial community and shifts the *Bacillus* community in the rhizosphere toward lower diversity and increased cooperativity. These new findings combined with the knowledge of relatedness, sociality, metabolic niche, and PGPR traits of *Bacillus* rhizosphere isolates, guided the design of plant beneficial *Bacillus* communities.

## RESULTS

### *B. velezensis* SQR9 alters the bacterial community in cucumber rhizospheres

*B. velezensis* SQR9 (hereafter referred to as SQR9) is a biostimulant that promotes the growth of cucumber (*Cucumis sativus*) through various plant growth promoting (PGP) mechanisms^27, 28, 32^. However, it is not known if and how this beneficial bacterium interacts with the resident rhizosphere community. To unravel this mystery, we grew cucumber seedlings with and without SQR9 in a pot experiment using sterile (SS) and natural soil (NS) as described in Fig. 1 (for details, see Methods section). As expected, inoculation with *B. velezensis* SQR9 (SS_SQR9 and NS_SQR9) significantly (*p* < 0.01) stimulated the growth of cucumber seedlings in both natural and sterile soil compared with the noninoculated control (Figs. S1A and S1B). Furthermore, the growth of cucumber plants in the sterile soil (SS treatment) was slightly slower than in the natural soil treatment (Fig. S1A and S1B).

**Fig. 1.**
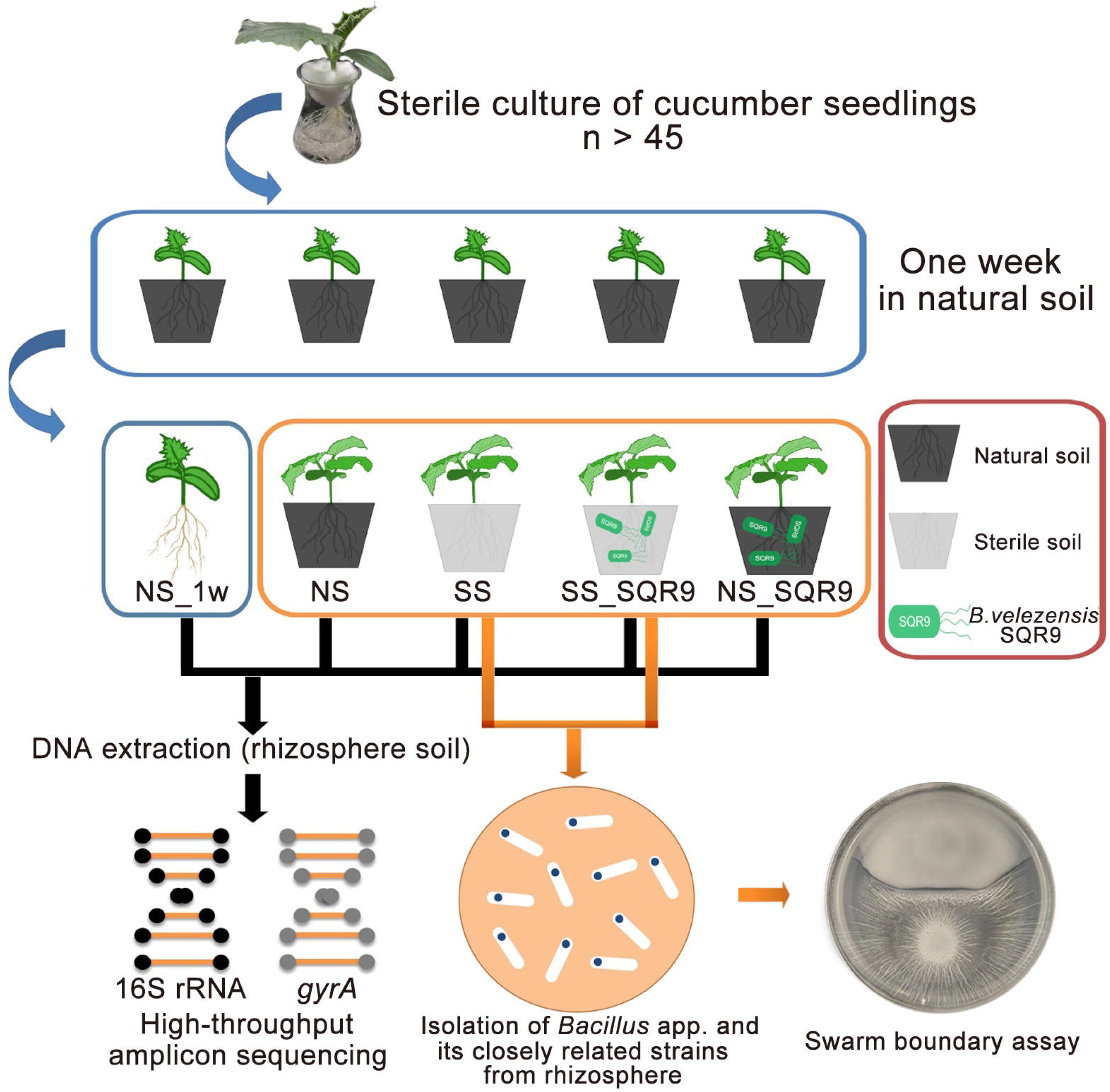
Overview of the experimental design and the effect of different treatments on cucumber plant growth. The workflow of the cucumber greenhouse experiment. Seedlings from the hydroponic culture system were transferred into fresh soil and incubated for one week. Then, DNA extracted from the rhizosphere soil of 9 out of 45 treatments (NS_1w) (blue panel) was used as a control. The remaining seedlings (orange panel) were transferred into fresh or sterile soil (NS, SS), with half of the soils inoculated with *B. velezensis* SQR9 (SS_SQR9 and NS_SQR9) and incubated for an additional two weeks. At this point DNA was extracted from all remaining rhizosphere soil experimental variants for 16S rRNA and *gyrA* amplicon sequencing. The rhizosphere soil from SS and SS_SQR9 variants (indicated by orange arrow) was also used for strain isolation. *Bacillus* isolates were tested for social interactions in a swarming assay.

To obtain detailed information on how strain SQR9 influences the resident bacterial community in the rhizosphere, we then used 16S rRNA amplicon sequencing, which provided comprehensive insight into the diversity of the bacterial community. In addition, we applied primers that amplify the gyrase-encoding gene *gyrA* of *Bacillus* species and relatives^33^. Compared to 16S rRNA-specific primers, the *gyrA* primers showed better phylogenetic resolution for the diversity of the *Bacillus* community^33^. By using both, more general 16S rRNA and more specific *gyrA* primers, we show here that the addition of SQR9 to sterile soil (SS_SQR9) significantly decreased the Shannon diversity index (H-index) of bacterial communities in the rhizosphere (*p* < 0.001) and *Bacillus* related communities (*p* < 0.001) (Figs. 2A and 2B). In contrast, in nonsterile soils (NS_SQR9) we did not detect a reduction in the H-index for the total bacteria but only for the *Bacillus* community and related species (Fig. 2A and 2B). Nonmetric multidimensional scaling (NMDS) analysis of 16S rRNA amplicons showed no significant separation of communities due to SQR9 treatment (Fig. 2C). In contrast, *gyrA* NMDS analysis confirmed the separation between SQR9-treated and untreated rhizosphere (Fig. 2D, Supplemental results 1). Overall, these results support the conclusion that SQR9 alters the composition of the indigenous bacterial community in the rhizosphere and that these shifts are more pronounced at lower taxonomic levels than at higher taxonomic levels, with more broadly distributed diversity.

**Fig. 2.**
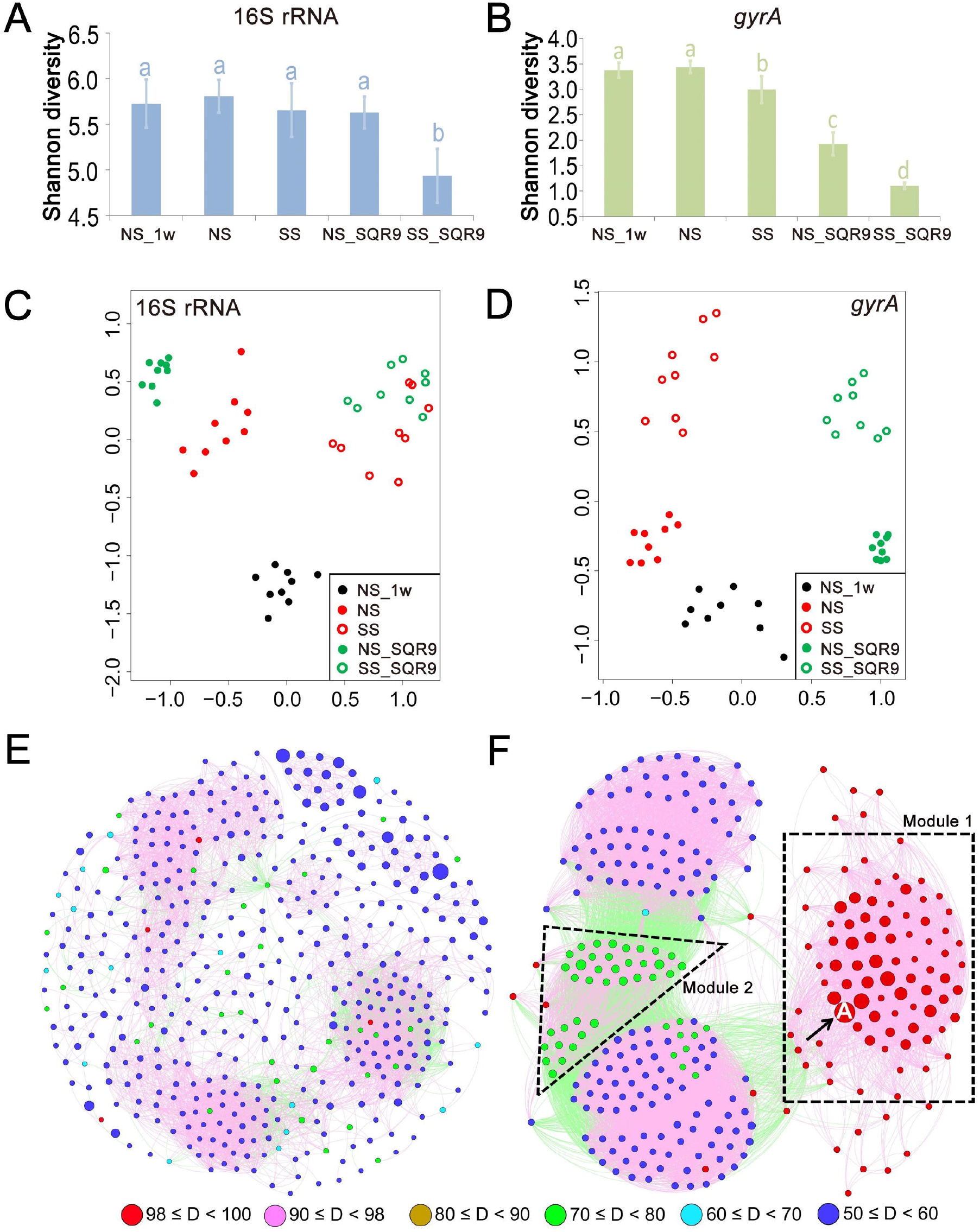
High throughput amplicon sequencing of 16S rRNA and *gyrA* in rhizosphere soil samples. (A) The Shannon diversity indices of the 16S rRNA sequence-based bacterial community. (B) The Shannon diversity indices of the *gyrA* sequence-based bacterial community. (C) Nonmetric multidimensional scaling plot of taxonomic similarity of the 16S rRNA gene (Bray-Curtis). (D) Nonmetric multidimensional scaling plot of the compositional similarity of the *gyrA* gene (Bray-Curtis). (E) The *gyrA* gene co-occurrence network of the non-SQR9-treated rhizosphere bacterial communities (NS and SS). (F) The *gyrA* gene co-occurrence network of the SQR9-treated rhizosphere bacterial communities (SS_SQR9 and NS_SQR9). Nodes with different colors depict the level of relatedness between *B. velezensis* SQR9 *gyrA* and other *gyrA* sequences. The capital letter A indicates the node of *B. velezensis* SQR9 in the community. Modules 1 and 2 show that the two groups of nodes gathered together in the community after inoculation with SQR9. For Figure E and F, red lines indicate positive correlations, and green lines indicate negative correlations.

Next, we applied network analyses of the *gyrA* gene co-occurrence patterns to further investigate the interactions between SQR9 and the indigenous rhizosphere bacterial community and their impact on community assembly. In the SQR9-treated rhizosphere, we observed a reduction in the total number of nodes and an increase in the number of links (including both positive and negative links) (Spearman’s correlation coefficient R > 0.80, *p* < 0.01, two-sided tests, Table S1). This result indicated that the addition of SQR9 strongly affects the cooccurrence of the indigenous rhizosphere bacterial community (Table S1, Fig. 2E, and 2F, Supplemental Results 2). Specifically, the color-coded nodes, which represent the degree of relatedness between the SQR9-*gyrA* and other *gyrA* sequences, confirmed a shift in community structure and enrichment of specific bacterial taxa in the rhizosphere (Fig. 2E and 2F). For example, in the untreated soil network, blue nodes representing phylogenetically distant members (50% ≤ D < 60%) dominated (Fig. 2E). In contrast, in the SQR9-treated soil, we observed enrichment of both the highly phylogenetically related (Module 1-red, 98% ≤ D < 100%) and moderately phylogenetically related members (Module 2-green, 70% ≤ D < 80%) with a concomitant increase in node connections (Fig. 2F). Moreover, the number of distantly related members (blue nodes) decreased in the SQR9-inoculated sample (Fig. 2F, Tables S2, Supplemental Results 2). These results reconfirm that SQR9 interacts with and shapes the structure of the *Bacillus* community and promotes the enrichment of more closely related members in the rhizosphere bacterial community.

### *B. velezensis* SQR9 enriches swarm merging interactions between isolates of the rhizosphere *Bacillus* sp. community

Given the results of bioinformatics analyses suggesting that SQR9 increases the connectedness of the *Bacillus* rhizosphere community, we set up experiments to test the hypothesis of their increased compatibility. Hence, we isolated spore-forming *Bacillus* strains and their relatives from the cucumber rhizosphere that had developed in sterile soil (SS) and sterile soil treated with SQR9 (SS_SQR9) (Fig. 1) and then tested their compatibility by a swarm encounter assay. When a boundary formed at the site of the swarm encounter, we assumed that the two strains were antagonistic or noncompatible, and if the two swarms merged, they were assumed to be compatible^24, 34^.

We obtained more than 200 spore-forming isolates and then selected 30 isolates from each treatment (SS and SS_SQR9) based on taxonomic criteria (*Bacillus* and related species), their swarming ability on 0.7% agar and their biofilm-forming activity (Supplemental methods 1). We then examined the phenotype of swarm interaction for 435 pairwise strain combinations (excluding self-self-pairs) from each treatment. Next, we compared the frequency of swarm phenotypes (merging, intermediate or boundary, Fig. 3A) from sterile untreated (SS) and SQR9-treated (SS_SQR9) rhizosphere soil and found that inoculation of the cucumber rhizosphere with SQR9 increased the frequency of the swarm merging phenotype of *Bacillus* sp. isolates. During swarming, only 29.7% of pairwise combinations of isolates obtained from sterile treatment (SS) merged, while 58.9% of pairs of isolates obtained from cucumber rhizospheres treated with SQR9 merged their swarms. Consistent with this, the frequency of *Bacillus* sp. boundary formation was lower in SQR9-treated soils (18.4% pairwise combinations) than in sterile soil (52.9%) (Fig. 3B, Table S3). These results are consistent with the bioinformatics data presented in Fig. 2E and 2F, and suggest that strain SQR9 alters the *Bacillus* community toward more compatible and potentially cooperative behavior.

**Fig. 3.**
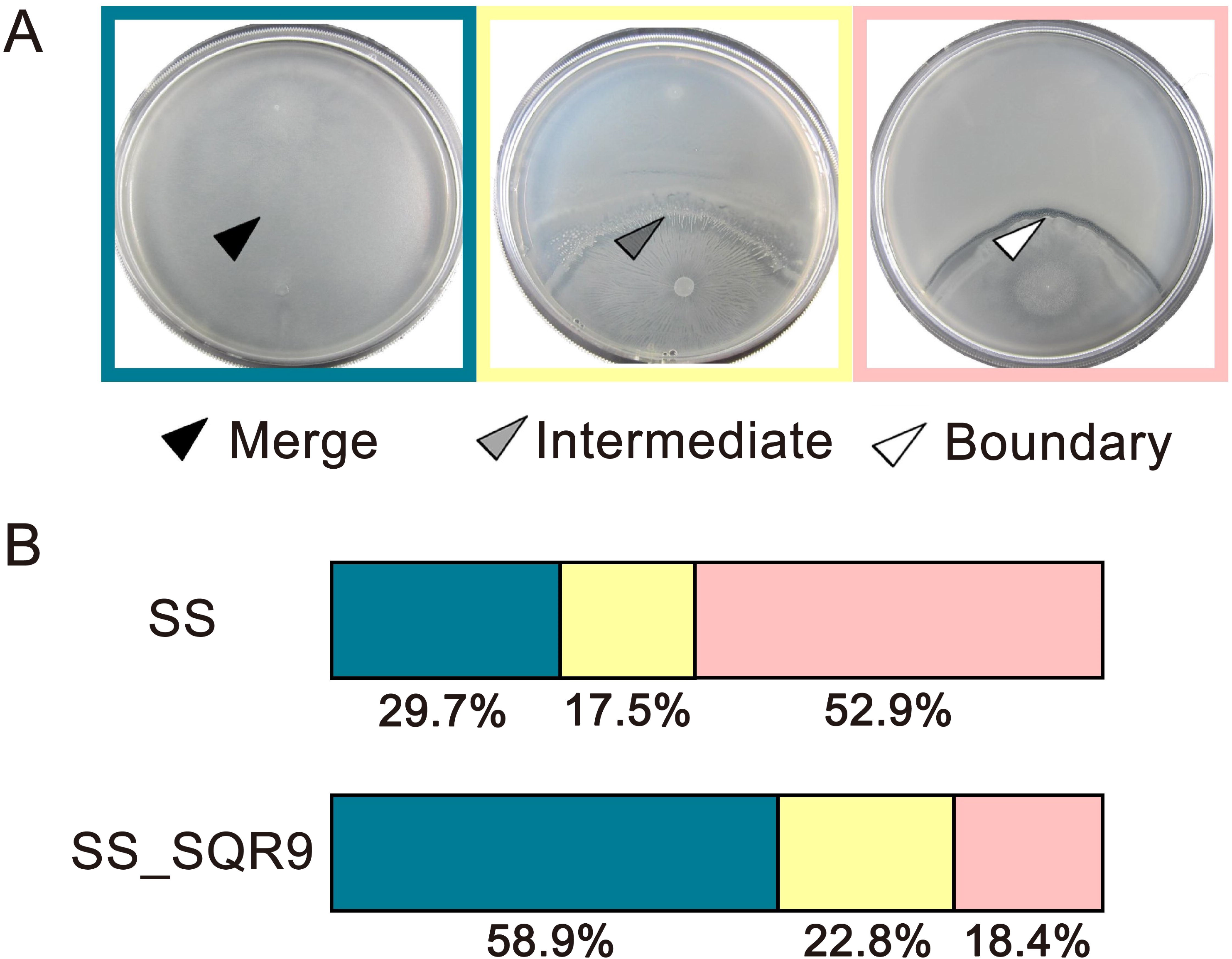
Soil inoculation with the strain *B. velezensis* SQR9 modifies the distribution of the pairwise swarm interaction phenotype between strains of *Bacillus* rhizosphere isolates. (A) Examples of merging or boundary formation between *Bacillus* strains isolated from the cucumber rhizosphere and staged separately on the agar surface of a 9-cm wide plate. Merging phenotype (black arrow); intermediate boundary (gray arrow), strong boundary (white arrow). The intermediate phenotype (gray mark) indicates a less striking but still visible line at the swarm encounter area. (B) The ratios of swarming interaction phenotypes (merging, intermediate and boundary) per 30 *Bacillus* isolates from the SS and SS_SQR9 treatments (see Table S3 for detailed distribution of swarm encounter phenotypes).

### Phylogeny and compatibility of rhizosphere isolates

To associate different swarming patterns from the SS and SS_SQR9 rhizosphere to the phylogenetic relatedness of interacting strains within each treatment, we determined the *gyrA* nucleotide identity between strains and constructed phylogenetic trees of isolated strains from treatments SS and SS_SQR9 (Fig. 4A and 4C, Supplemental Results 3), which are referred to as isolated communities. Strains isolated from the SS treatment in most cases showed more than 90% identity at the *gyrA* gene and a lower frequency of the swarm merging phenotype (Fig. 4B). In contrast, the SS_SQR9 treatment strains were on average less related (down to 84%) but more compatible with a higher frequency of the merging phenotype (Fig. 4D). Merging was a predominant phenotype between SS_SQR9 treatment strains, with *gyrA* identity ranging from 96 to 99.5% (Fig. 4C and D).

**Fig. 4.**
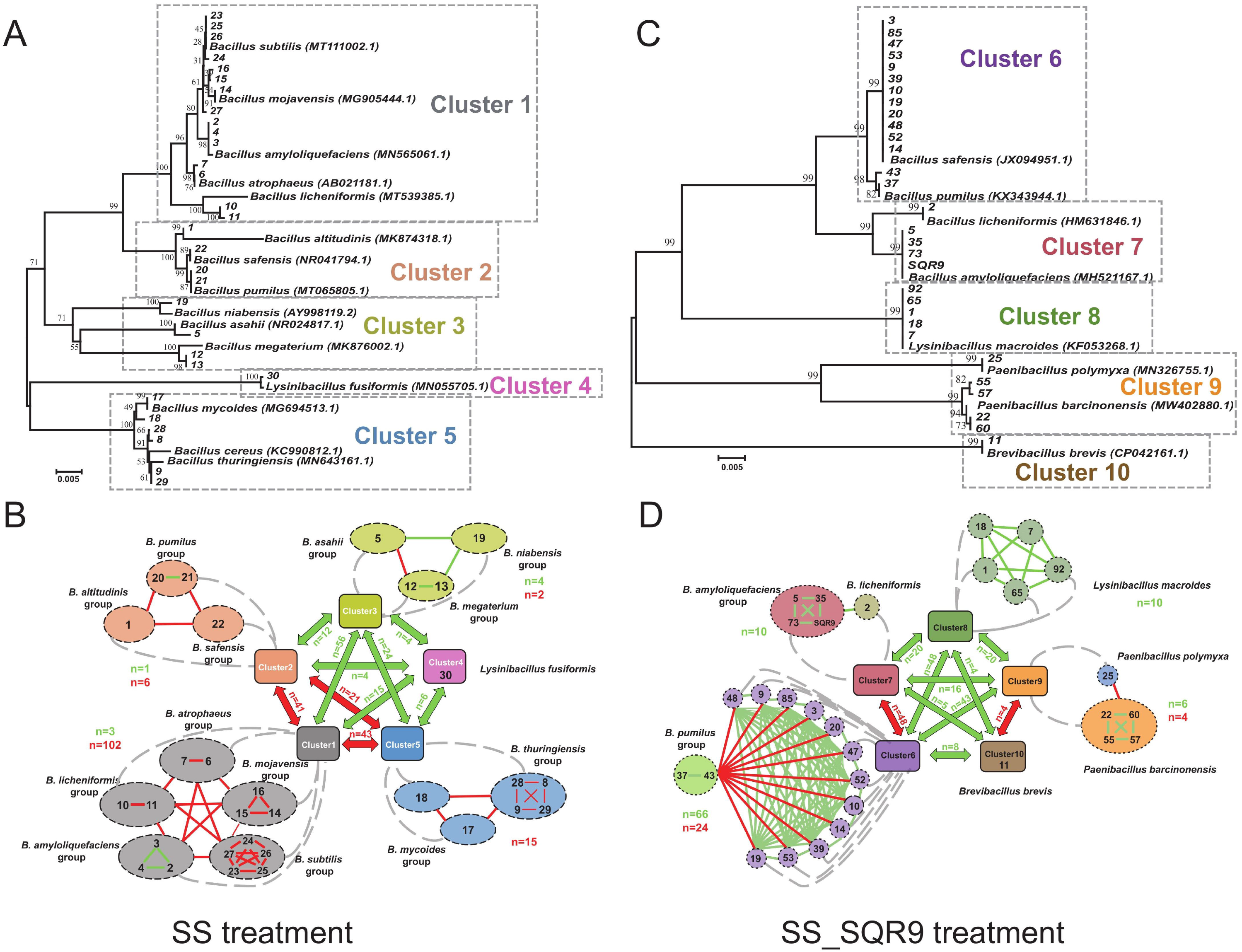
Recognition groups of the cucumber rhizosphere *Bacillus* population isolated from the SS and SS_SQR9 treatments. (A) The minimum-evolutionary tree based on full-length *gyrA* gene sequences from 30 strains isolated from the cucumber rhizosphere - SS treatment. (B) Swarm interaction network of the strains isolated from the cucumber rhizosphere in the SS treatment. (C) The minimum-evolutionary tree based on full-length *gryA* gene sequences from the 30 strains isolated from the cucumber rhizosphere in the SS_SQR9 treatment. (D) Swarm interaction network of the strains isolated from the cucumber rhizosphere in the SS_SQR9 treatment. Green connection lines represent strains with a merging phenotype; red connection lines represent boundary formation. Colors depict different groups, and n represents the number of pairwise combinations displaying the swarm interaction phenotype within a group or between clusters (the numbers in green indicate the number of merging phenotypes, and the numbers in red indicate the number of boundary phenotypes).

For strains in the SS treatment, the boundary phenotype was predominant within the arbitrary clusters (boundaries (B)=125, merging (M)=8), with merging and boundary formation between clusters present almost equally (B=105, M=121) (Fig. 4B). Among strains isolated from the SQR9-treated rhizosphere soil (SS_SQR9), the merging phenotype dominated within species clusters and between arbitrary clusters (B=28, M =92). Although some strains of two closely related species within arbitrary clusters formed boundaries (e.g. *P. polymyxa* and *Paenibacillus* or *B.safensis* and *B. pumilus*), some also merged (e.g., *B. licheniformis* and *B. amyloliquefaciens*). Moreover, we observed an enrichment of merging between strains from different arbitrary clusters (B=52, M=164) (Fig. 4D). Overall, and in line with our bioinformatics data (Fig. 2), these observations suggest that SQR9 rhizosphere inoculation reduces the frequency of antagonists and makes the rhizosphere bacterial community more compatible.

### Learning from nature: SQR9 enrichment of moderately related swarm mergers promotes plant growth

Thus far, our results have shown that *B. velezensis* SQR9 added to the rhizosphere changes the composition and social interactions of the bacterial community. However, how this information can be used to improve the design of PGP consortia has yet to be determined. It has been shown previously that highly related swarm compatible strains coexist on plant roots^34^ and that moderately related strains also merge swarms^23^. Hence, we reasoned that moderately related compatible strains will coexist on plant roots and exhibit less resource competition than closely related strains, which will improve the activity of moderately related PGP consortia. To test this hypothesis, we first determined the resource competition among the candidate PGPR strains in our consortia. We measured the carbon source utilization of 30 strains from the rhizosphere treated with SQR9 using the GEN III MicroPlate test assay performed by the Biolog system. Principal component analysis (PCA) showed that the patterns of carbon source utilization correlated strongly with the phylogenetic relatedness of the *Bacillus* isolates (Fig. 5A and 4C). In conjunction with information on swarming patterns and relatedness of strains to SQR9 (Fig. S2), this information provided the basis for consortia design.

**Fig. 5.**
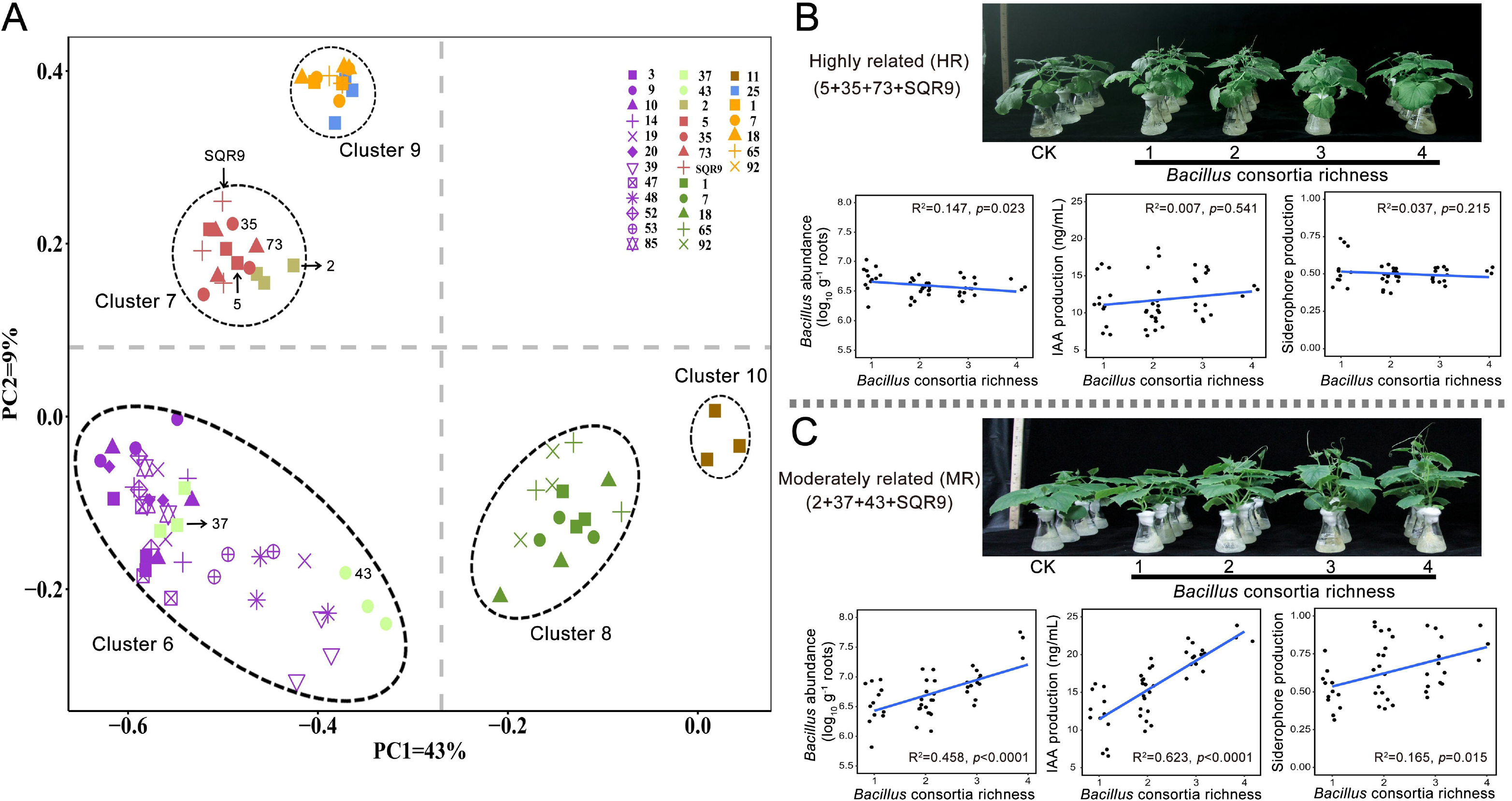
Carbon resource competition and sociality are two important parameters for the smart design of PGP *Bacillus* consortia. (A) Principal component analysis (PCA) of the carbon source utilization pattern on Biolog GENIII plates of 30 strains isolated from the cucumber rhizosphere (SS_SQR9 treatment). The plotted data are averages of three independent experiments. The locations of *Bacillus* isolates used to build consortia are marked. Briefly, *Bacillus velezensis* 5, 35, 73 and SQR9 were used to build highly related (HR) consortia, and *Bacillus licheniformis* (2), *Bacillus pumilus* (37 and 43) and *Bacillus velezensis* (SQR9) were used to build moderately related (MR) consortia. (B) Increasing the richness of highly related (HR) consortia (5+35+73+SQR9) had no effect on cucumber growth, colonization abundance, IAA and siderophore production under laboratory conditions. (C) In contrast, increasing the richness of moderately related (MR) consortia (2+37+43+SQR9) stepwise improved cucumber growth, colonization abundance, IAA and siderophore production under laboratory conditions.

As the final objective was to enhance plant growth, we applied the insights gained to mimic the consequences of SQR9 soil inoculation in the rhizosphere. We hypothesized that phylogenetic relatedness-based sociality and competition for carbon resources represent fundamental knowledge for the rational design of PGP consortia. To test this prediction, we compared two sets of consortia: the highly related swarming consortia (HR, 100% identity of the *gyrA* gene, isolates 5, 35, 73, SQR9) and the moderately related swarming consortia (MR, 70% ≤ D < 80%, isolates 2, 37, 43, SQR9) (strain traits indicated in Fig. 5A, 4C and S2). As it has been previously shown that increasing the richness of PGP consortia also positively affects PGP activity on tomato plants^35^, we tested the effect of MR and HR consortia with increasing richness on cucumber growth in an experimental hydroponic system and potting experiments with natural soil.

As predicted, we did not detect an increase in PGP activity, an improvement in cucumber root colonization or an increase in IAA and siderophore production by combining HR strains, regardless of whether we used one or multiple strain consortia (Fig. 5B, S3A and S3B). MR consortia, on the other hand, resulted in an increase in root colonization and IAA production with increased MR richness (R^2^ = 0.458, *p* < 0.0001; R^2^=0.623, *p* = 0.0001) (Fig. 5C). Concerning siderophore production, the effect was significant (*p* < 0.15), but the correlation with MR richness was less strong (R^2^ = 0.165) (Fig. 5C). Additionally, the MR consortia improved cucumber growth in hydroponic (Fig. S3C and S3D) and potting experiments with nonsterile natural soil (Fig. S4A), again with significant increases in shoot height and dry shoot weight with increasing richness (Fig. S4B and S4C). Although individual MR isolates showed low PGP potential (Fig. S5), their ability to promote plant growth increased when multiple strains were combined (Fig. 5C, Supplemental Results 4). Our results indicate that mixing moderately related swarm-merging strains significantly promotes PGP activity in a richness-dependent manner, highlighting the importance of relatedness-dependent ecological compatibility and niche breadth for consortia design.

Next, we constructed two types of consortia (HR and MR) with increasing richness to test our prediction further in a hydroponic system. Specifically, each consortium contained a mixture of one to eight strains from our collection of 60 cucumber rhizosphere strains. These gave 150 combinations for each consortia type: 60 with a single strain, 30 with two strains, 30 with four strains, and 30 with eight strains (Table S4). The HR consortia contained selected strains that exhibited a merging phenotype between their swarms and high competition for carbon sources (Table S4); the MR consortia contained selected strains exhibiting a merging phenotype between their swarms and lower competition for carbon sources (Table S4). We hypothesized that the HR and MR consortia would affect strain abundance, IAA production and siderophore production differently and that MR consortia would have more plant growth beneficial properties. Consistent with the trends shown in Fig. 5, in the hydroponic experimental system, the MR consortia showed improved performance in the hydroponic system in terms of PGP activity (*Bacillus* abundance, IAA production; siderophore production), which was evident with increasing richness (Fig. 6). Although in experiments with eight different strains tested in 150 combinations, the scattering of results was too high for us to predict significant correlations, the trend of increasing PGP activity with increasing richness was clear (Fig. 6). These richness dependent effects were not observed for HR consortia. We concluded that the positive effect of MR consortia richness on PGP activity may be due to low competition among individuals and the potential synergies created by complementary niches and thus more effective use of available resources (Fig. 6).

**Fig. 6.**
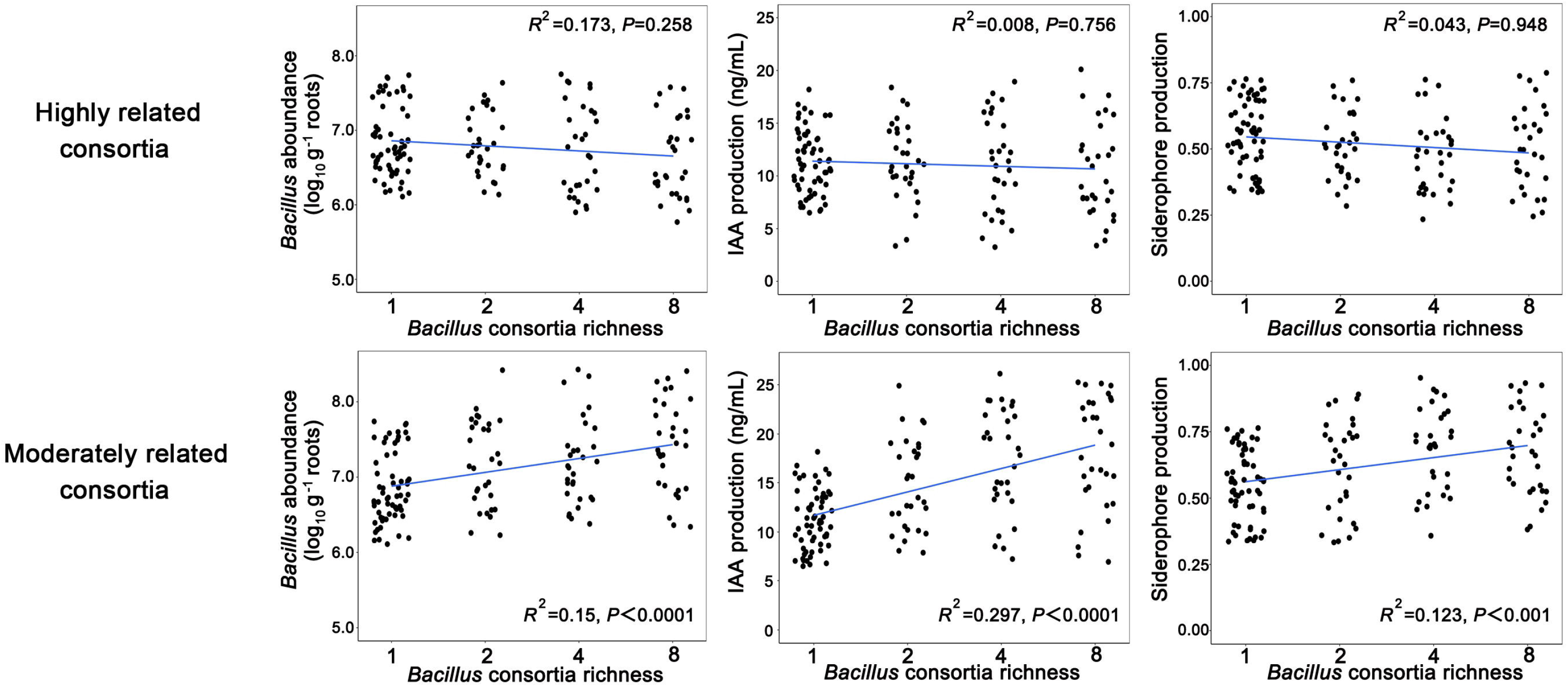
Comparison of the PGP properties (colonization abundance, IAA and siderophore production) of two types of consortia reassembled from *Bacillus* isolated from cucumber rhizosphere soil. A synergistic effect was observed in moderately related (MR) consortia; increasing the richness of moderately related (MR) consortia stepwise improved colonization abundance, IAA and siderophore production.

## DISCUSSION

Plant-associated beneficial microorganisms show great promise for improving crop quality and productivity^36, 37^. However, their use is hampered by our limited understanding of the relationships between the effects of microbial inoculants on the plant and its rhizosphere microbiome^37^. Here, we provide evidence that *Bacillus velezensis* SQR9 changes the composition of indigenous rhizosphere bacteria, and *Bacillus* species in particular, toward a less competitive and more cooperative community in which highly and moderately related strains are enriched (Fig. 7). Furthermore, our results provide the first evidence that the associated sociality shifts community-level functionality, as reflected in the improved PGP activities of moderately related consortia, which is important for developing more reliable PGP inoculants.

**Fig. 7.**
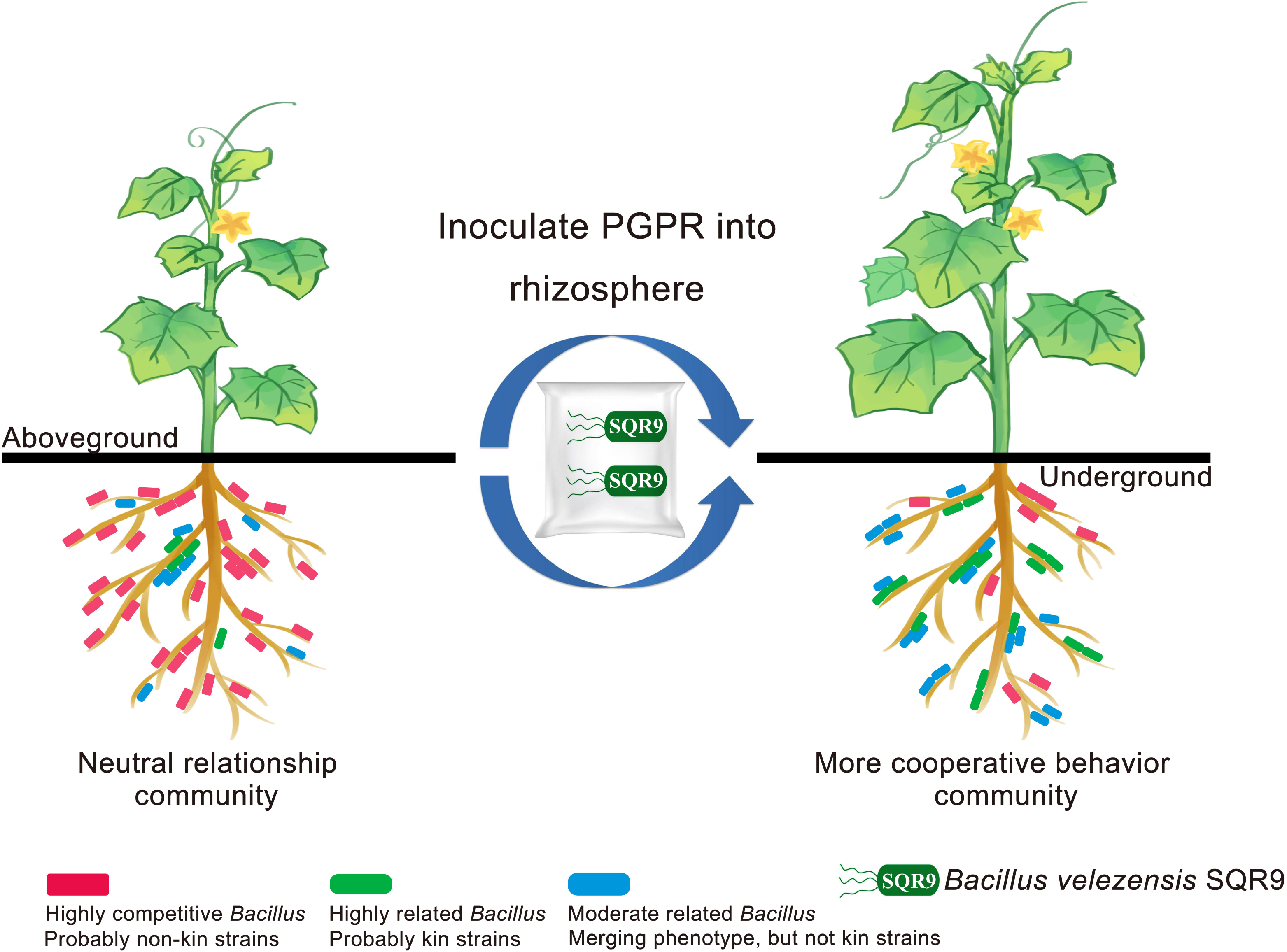
New insight and schematics of the ecological impact of microbial inoculants (*B. velezensis* SQR9) on *Bacillus* and its closely related community in the rhizosphere. Strain SQR9 alters the *Bacillus* community toward more cooperative and compatible behavior in the cucumber rhizosphere, leading to the promotion of plant growth.

### The application of SQR9 alters the composition of the *Bacillus* community and related species in the rhizosphere

Although *Bacillus* species are known for their PGP activities and are commonly applied in agroecosystems to improve plant health^38, 39^, their impact on the rhizosphere bacterial community is still poorly understood. Previous studies examining the effects of inoculants, such as *B. amyloliquefaciens* FZB42, BNM122 and *B. subtilis* PTS-394, by amplicon sequencing of 16S rRNA genes did not detect a significant impact on the overall bacterial community composition^40–42^. However, in this study, we complemented 16S rRNA gene analysis with *gyrA* amplicon sequencing. Although we also detected a decrease in 16S rRNA diversity in the SQR9-treated SS-rhizosphere, the *gyrA* targeting approach provided a means to unravel previously undetected dramatic changes in the composition of *Bacillus* species and closely related genera (*Paenibacillus, Brevibacillus, Lysinibacillus*), emphasizing the importance of intra- and interspecific competition. This approach required the development of a new tool – the *gyrA3* primer pair, which covers the broad diversity of *Bacillus* species and relatives^33^. In this work, we provide evidence that this primer amplifies *gyrA* genes of a rhizosphere bacterial community associated with an agriculturally important plant and that by targeting lower taxonomic units one can improve the detection and resolution of the rhizosphere microbiome dynamics by amplicon sequencing. Additionally, this work shows that SQR9 causes a dramatic shift toward a less diverse community with a higher proportion of highly interconnected strains that increased in frequency and were either highly related (98-100% identity, red group) or moderately related (70-80% identity, green group in Fig. 2F). In contrast, the genotypes with 50-60% *gyrA* identity (blue group) decreased in frequency. SQR9 also increased the number of connections within each group, revealing a positive correlation among the close relationship nodes (Fig. 2E and 2F).

### Strain SQR9 changes social interactions in the rhizosphere bacterial community

Although the importance of the rhizosphere microbiome to plant health^43^ and the potential benefits of bioinoculants are widely recognized^5, 36^, the effects of bioinoculants on bacterial social interactions in the rhizosphere have been largely ignored. This work shows that strain SQR9 causes a dramatic shift in the community, with a higher proportion of strains that are related and thus more interconnected with SQR9 (Fig. 2E and 2F). Furthermore, compatibility among isolates from the rhizosphere treated with SQR9 dramatically increased (Fig. 3B). Moreover, the patterns of the *gyrA* gene co-occurrence and the results of swarming assays provided the first evidence that *Bacillus* and its closely related species are important targets of SQR9 activity in the rhizosphere (Fig. 2E, 2F and S2, Table S2).

What could be the mechanisms behind these effects? It seems that strain SQR9 outcompetes closely and more distantly related species. For example, the untreated isolated rhizosphere community comprised 15 *Bacillus* species and only one isolate of *Lysinibacillus* (Fig. 4A). In contrast, the SQR9-treated isolated community comprised four *Bacillus* species and three more moderately related species (Fig. 4C), confirming the strong antagonism of SQR9 against closely related species in the rhizosphere. In fact, *B. velezensis* strains, including SQR9, are known to produce a diverse range of bioactive secondary metabolites^44–47^. Therefore, intense competition may lead to niche emptying and consequently provide an opportunity for compatible strains to grow, which is confirmed by our results.

The enrichment of compatible strains (Fig. 3) can also be explained by kin discrimination (KD)^20^. KD is mediated in *Bacillus* species by mechanisms involving intercellular attack and defense molecules with extensive variation in genes and unique combinations in different strains^23^. Antagonistic actions can be identified by the appearance of visible boundaries (boundaries, clearing zones) between swarms^25^, with strains forming a boundary on semisolid media unable to coexist on plant roots, while swarm mergers join to form a plant root biofilm. Killing competitors can be advantageous, and indeed, we observed that strain SQR9 promoted the enrichment of compatible strains within specific clusters (Fig. S2) and reduced diversity within species clusters (compare Fig. 4A and 4B). Although swarm merging has been generally associated with *Bacillus* intraspecific sociality (including all intraspecies references) and some more distantly related *Bacillus* species^23^, we found that moderately related species in the rhizosphere, including *Bacillus, Paenibacillus* and *Lysinibacillus* species, can also merge their swarms.

By looking at the community from two perspectives, first, how the addition of strain SQR9 changes community relatedness and compatibility in relation to SQR9, and second, how the strains in the rhizosphere community interact with each other after SQR9 inoculation (Figure 4), we conclude that strain SQR9 shifts the community toward cooperativity. Overall, our results demonstrate that SQR9 via antagonism provided an opportunity for compatible strains to grow and consequently decrease competition. This is consistent with the theory that a certain level of competition is particularly important for the development of cooperation in bacteria^18, 48^.

### Smart design of synthetic *Bacillus* consortia with defined ecological (social) interactions

Plant-associated microbial communities have numerous potential applications in biotechnology, especially in agriculture^49^. Recently, microbial consortia with lower complexity have been studied and used as model systems for the controlled assessment of ecological, structural, and functional properties of microbial communities^50^. However, a major challenge is to rationally engineer beneficial consortia with robust survivability and activity for functionally reliable applications under field conditions^51^. Addressing this challenge requires the integration of multiple approaches, including the reverse engineering of natural communities (e.g., inference-based co-occurrence analysis) and the rational engineering of beneficial consortia with desirable interactions for community-level functionality and robustness^52^. Despite the fact that several underlying factors could directly or indirectly affect the performance of synthetic *Bacillus* consortia in the rhizosphere^53^, we show that sociality and competition for carbon among these strains are important parameters that should be considered in the development of efficient plant probiotics (PGP inoculants). The results show that mixing compatible strains that do not compete for the same resources leads to more efficient PGP inoculants, and we confirmed their PGP activity in both hydroponic and natural soil systems (Fig. 5C and S4).

Overall, our work provides a general ecological framework for the intelligent assembly of *Bacillus* consortia for more efficient and reliable applications. First, the work provides evidence that the application of the beneficial strain SQR9 shifts the bacterial community in the rhizosphere and, in particular the *Bacillus* community toward increased cooperativity. Second, the results suggest that by including the ecological mechanisms (sociality and competition for carbon) used by microbial inoculants, we can improve guidelines and develop more effective products for sustainable agriculture.

## MATERIALS AND METHODS

### Bacterial strains, isolation and culture conditions

We used *Bacillus velezensis* SQR9 (CGMCC accession no. 5808, China General Microbiology Culture Collection Center) as our model strain to assess its ecological impact on the resident bacterial community in the rhizosphere. Additional natural isolates of *Bacillus* and other spore formers were obtained from two kinds of cucumber rhizosphere soils, where cucumber seedlings grown in natural soil were transferred to sterile soil only (SS) or additionally inoculated with SQR9 (SS_SQR9) (see experimental design Figure 1). Two hundred spore-forming isolates were obtained by heat treatment of rhizosphere soil samples and then 30 representative isolates for both SS and SS_SQR9 treatments were selected for further experiments based on the following criteria: a, three metabolic tests (the catalase test, the Voges-Proskauer test and anaerobic growth on agar) and the 16S rRNA nucleotide identity identified them as *Bacillus*^54^; b, the ability to form pellicles (floating biofilm) in MSgg medium; and c, the ability to swarm on 0.7% agar. *Bacillus* strains were routinely grown in LB medium and, where appropriate, in B medium prepared as described^34^.

### Experimental design

To test whether the application of SQR9 to the rhizosphere shapes the overall bacterial community structure, we designed a cucumber greenhouse experiment, as presented in Figure 1. Two-week-old cucumber seedlings from a hydroponic culture system^55^ were transferred to pots filled with 200 g natural soil and incubated for one week, and then DNA from rhizosphere soil was extracted from 9 out of 45 treatments (NS_1w) (blue panel, Fig. 1). The remaining seedlings (orange panel, Fig. 1) were transferred to pots filled with 200 g natural or sterile soil (NS, SS) or both types of soils inoculated with 10 mL suspensions (10^7^ CFU mL^−1^) of *B. velezensis* SQR9 (NS_SQR9 and SS_SQR9) and incubated for an additional two weeks. At this point, DNA was extracted from the rhizosphere soil for 16S rRNA and *gyrA* amplicon sequencing. The rhizosphere soil from SS and SS_SQR9 variants (indicated by the orange arrow, Fig. 1) was also used for *Bacillus* spp. strain isolation. Strains were tested for social interactions in a swarming assay (Fig. 1) as described below. Plant shoot height and shoot dry weight were measured after four weeks of incubation.

### DNA extraction and sequencing

Cucumber rhizosphere soils were collected as described by Chaparro et al.^56^. Briefly, cucumber roots were gently removed from the soil, and the loose soil (> 1 mm, not the rhizosphere soil) was manually removed with sterile rubber gloves, leaving approximately 1 mm of soil on the roots. Rhizosphere soils were collected, and total DNA was extracted from 0.25 g of rhizosphere soil using the PowerSoil DNA Isolation Kit (Mo Bio Laboratories, Inc., Carlsbad, CA, USA). To minimize DNA extraction bias, successive DNA extractions of each sample were pooled before performing polymerase chain reaction (PCR). A NanoDrop ND-2000 spectrophotometer (NanoDrop, ND-2000, Thermo Scientific, 111 Wilmington, DE, USA) was used to assess DNA quality according to the 260/280 nm and 260/230 nm absorbance ratios^57^.

Amplification of the V3-V4 hypervariable region of the bacterial 16S rRNA gene was performed to assess the bacterial community using the primers 338F: 5’-CCTACGGRRBGCASCAGKVRVGAAT-3’ and 806R: 5’-GGACTACNVGGGTWTCTAATCC-3’. For *Bacillus* and its close relatives, amplification of the *gyrA* housekeeping gene was performed using the primers 243F: 5’-GCDGCHGCNATGCGTTAYACTC-3’ and 736R: 5’-CGGACAAGMTCWGCKATTTTTTC-3’ to assess the community composition. Nine 16S rRNA samples and three *gyrA* gene samples were sequenced per treatment. In comparison to 16S rRNA, these primers target *Bacillus* species, and show better phylogenetic resolution for *Bacillus* community diversity than 16S rRNA primers^33^. PCR amplifications were combined in equimolar ratios and sequenced on an Illumina MiSeq instrument.

### Bioinformatics analysis

The relative abundance of all *gyrA* genes was used for network analysis. The network analysis was performed using the Molecular Ecological Network Analyses Pipeline (MENAP) (http://ieg2.ou.edu/MENA/main.cgi)^58^. According to the pipeline, network analysis contains two steps: network construction and network analysis. The network construction includes data updates, data standardization, pairwise similarity of relative abundance across different samples and determination of the adjacency matrix by an RMT-based approach. The network analysis includes network module detection, network overall topological structure, and topological role identification of individual nodes^59^. The Shannon diversity index calculation and Bray–Curtis dissimilarity-based NMDS analysis were performed based on the rarefied table of the sequencing data using the vegan R package (v.2.5–2) (https://cran.r-project.org/package=vegan).

### Swarm boundary assay

To test the social interaction between approaching swarms of different *Bacillus* spp. isolates, 9 cm plates containing B-medium with 0.7% agar were prepared fresh. *Bacillus* spp. strains were grown on solid Luria-Bertani plates at 30 °C for 16 h. Cells were transferred into 3 mL of liquid B-medium and shaken overnight at 30 °C. Overnight cultures were then diluted to an optical density (OD_600_) of 0.5, and 2 μL was spotted on the plates at each side of the agar plate. The plates were dried in a laminar flow hood for 20 min, incubated for 2 days at 30 °C, and photographed. Three phenotypes (merging, intermediate and boundary) were assigned to 870 pairs of swarms as described previously^34^.

### Establishing a *Bacillus* community richness gradient for two kinds of consortia to test the effects of PGP *in vitro* and greenhouse experiments

A probiotic *Bacillus* community richness gradient was created by establishing four richness levels (1, 2, 3 and 4) using *Bacillus* strains isolated from rhizosphere soil inoculated with SQR9 (SS_SQR9). We used a substitutive design where the total probiotic community biomass was kept constant at all richness levels. The same experimental design was followed in both the greenhouse and *in vitro* experiments. For further experiments, a representative *Bacillus* community for highly related (HR) consortia was built by using our collections with designated numbers: *Bacillus velezensis* (5), *Bacillus velezensis* (35), *Bacillus velezensis* (73), and *Bacillus velezensis* (SQR9); a representative *Bacillus* community for moderately related (MR) consortia was built by using isolates *Bacillus licheniformis* (2), *Bacillus pumilus* (37), *Bacillus pumilus* (43), and *Bacillus velezensis* (SQR9). These two kinds of consortia with varying richness were used to test the PGP effects *in vitro* and in greenhouse experiments.

To test the PGP effects of different *Bacillus* consortia *in vivo*, two forms of greenhouse experiments were performed. (1) The first was a hydroponic culture system, where we tested consortia with 4 *Bacillus* strains. The planting system was carried out in an aseptic conical flask containing 50 mL ¼ sterile Murashige Skoog (MS) medium, as described by Qiu et al.^60^. Briefly, cucumber seeds of the cultivar Jinchun 4 were surface disinfected in 2% NaClO solution for 15 min, washed thoroughly with distilled sterile water and planted in sterile vermiculite in a tissue-culture container. The seeds were allowed to germinate and grow for approximately 4 days in a growth chamber at 23 °C with a 16 h light and 8 h dark photoperiod. When two cotyledons appeared, the seedlings were individually transplanted into an aseptic conical flask containing 50 mL ¼ sterile MS medium. After 4 days of growth, plants were inoculated to a final OD_600_=0.02 with the *Bacillus* community grown for 4 h and placed on an orbital shaker at 50 rpm in the growth chamber at 70% humidity with natural light and 28±2 °C during the day and 22±2 °C at night for another two weeks. Each conical flask was treated as one biological replicate, with nine replicates set up for each treatment and nine replicates for the noninoculated control. Plant shoot height and shoot dry weight were measured after three weeks of incubation. (2) The second was a natural soil system, where the plant growth-promoting efficiency of each *Bacillus* community (composed of one to eight strains) was assessed in a greenhouse assay with natural, nonsterile soil. The trial was conducted from August 17 to November 8 in 2019 in the greenhouse at Nanjing Agricultural University. The soils used for the greenhouse assay were collected from a field site with a history of cucumber cultivation. The field site was located in Nanjing, Jiangsu Province, China and the soil had the following properties: pH 6.4, organic matter 18.6 g kg^−1^, available N 121 mg kg^−1^, available P 56 mg kg^−1^, and available K 89 mg kg^−1^. Two-week-old cucumber seedlings were transplanted to seedling pots with 5 kg of soil. After 7 days of growth, plants were inoculated with assembled consortia by drenching the pots to a final concentration of 10^7^ CFU g^−1^ soil. Each treatment was replicated 18 times, and the experimental plan included three blocks in a completed randomized design (9 plants for each block). The pots were incubated in a growth chamber at 70% humidity with natural light and 28±2 °C during the day and 22±2 °C at night. Plants were irrigated regularly with ½ Hoagland medium as described by Qiu et al.^60^. Nine randomly selected plants for each treatment were harvested after 55 days. Plant shoot height and shoot dry weight were measured.

### Root colonization assay

Two-week-old cucumber seedlings were transferred to 30 mL of 1/4 Murashige and Skoog (MS) culture medium in an aseptic conical flask. The medium hosting the plant was then inoculated with *Bacillus* spp. strains at the final OD_600_=0.02 in medium (approximately 1 × 10^6^ CFU mL^−1^), and placed on an orbital shaker at 100 rpm in the growth chamber at 30 °C. After 2 days, the bacterial cells that colonized the cucumber root surfaces were collected and quantified using the method described by Qiu et al.^61^. Briefly, cucumber roots with colonized cells were aseptically removed from the container, washed three times in sterile water, and placed on axenic filter paper to remove the planktonic cells. Then, the roots were placed into a 250 mL sterile flask containing 45 g of glass beads (6 mm in diameter) and 100 mL of sterile water, which was shaken vigorously with a vortex mixer for 10 min to separate the attached cells from the roots.

### Carbon source utilization assay

The carbon source utilization patterns of 60 *Bacillus* isolates from cucumber rhizosphere soils were recorded by the GEN III MicroPlate test assay performed with the Biolog system (Biolog, CA, USA). Briefly, bacterial suspensions were prepared and inoculated in a Biolog GEN III MicroPlate according to the manufacturer’s instructions (http://www.biolog.com). The utilization of carbon sources was determined after 12 h using Protocol A of Biolog’s MicroStation™ System^62^.

### Multiple combinations analysis the correlation between social interactions among community members and PGP phenotypic characterization

We assessed consortia functionality using the same methods as for the single isolates. In this experiment, we focused on three PGP phenotypic traits: colonization abundance and IAA and siderophore production. Bacteria from frozen stock were pregrown overnight in liquid LB at 30 °C, pelleted by centrifugation (4,000 ×g, 3 min), washed three times in 0.85% NaCl, and the cell density was adjusted to an OD_600_ of 1.0 prior to further experiments. We then built *Bacillus* consortia using a substitutive design with richness levels of 1, 2, 4, and 8 strains from the collection of 60 *Bacillus* isolates from the cucumber rhizosphere and tested their effect in natural soil system. There were 150 combinations with each consortia type, 60 with a single strain, 30 with two strains, 30 with four strains, and 30 with eight strains (Table S4). The consortia design was conceived to ensure that each isolate was present at comparable frequency at each diversity level, allowing for separation of the effects of bacterial richness and composition^35^. Detailed information on the components of the two kinds of *Bacillus* consortia is shown in Table S4, and the selection criteria were as follows: for HR consortia, we selected strains with a merging phenotype and high carbon resource competition; for MR consortia, we selected strains with a merging phenotype and lower carbon resource competition. *Bacillus* consortia were grown under the same conditions as the single isolates, after which the different traits were measured at the community level.

### Statistical analysis

Figures were produced using the GraphPad Prism 8 or ggplot2 R package. Detailed statistical analysis is described in the figure legends. The data on all individual *Bacillus in vitro* performances (five PGP traits) were standardized between 0 (minimum value across all treatments) and 1 (maximum value across all treatments) and used in subsequent calculations and analysis as described above.

### Data availability

Amplicon sequencing reads from the 16S rRNA gene and *gyrA* gene are available at NCBI Sequence Read Archive under accession number PRJNA879238.

## Supporting information

Supplemental Figure 4A_distance

Supplemental Figure 4C_distance

Supplemental results and methods

## AUTHOR CONTRIBUTIONS

ZX, IMM designed the study, YL, TY performed the experiments. YL, ZX, WX, YM, PS, IMM analyzed the data. ZX, WX created the figures. ZX wrote the first draft of the manuscript, ZX, IMM, PS, NZ, RZ and QS revised the manuscript.

## ACKNOWLEDGMENTS

This work was financially supported by the National Natural Science Foundation of China (31972512, 42090064 and 32072675), the National Key Research and Development Program (2021YFD1900300), the National Scientific and Technological Program on Basic Resources Investigation (No. 2019FY102000) and Slovenian Research Agency national program grant P4-0116 and research projects J4-9302 and J4-8228.

---

Fig. S1 Inoculation with *B. velezensis* SQR9 significantly stimulated cucumber growth in both natural and sterile soils compared with the noninoculated control (NS, SS). (A) Inoculation with SQR9 significantly increased cucumber shoot height. (B) Inoculation with SQR9 significantly increased cucumber shoot dry weight. Different letters above the bars indicate significant differences between all experimental variants on the plot (*p* < 0.01, *t test*, n=9, mean ± standard).

Fig. S2 The minimum-evolutionary tree based on full-length *gyrA* gene sequences from the 30 strains isolated from the cucumber rhizosphere in the SQR9 treatment. For each isolate, the similarity of the *gyrA* gene to SQR9 and the swarm interaction phenotype with SQR9 are also presented in the figure.

Fig. S3 Positive effect of moderately related (MR) consortia (2+37+43+SQR9) strain richness on cucumber growth. Increasing the richness of highly related (HR) consortia (5+35+73+SQR9) had no effect on cucumber shoot height (A) and shoot dry weight (B) in the sterile hydroponic culture system. In contrast, increasing the richness of moderately related (MR) consortia (2+37+43+SQR9) stepwise improved the cucumber shoot height (C) and shoot dry weight (D) under the same conditions. Different letters above the bars indicate significant differences (*p* < 0.01, *t test*, n=9, mean ± standard deviation).

Fig. S4 Increasing the richness of moderately related (MR) consortia (2+37+43+SQR9) significantly improved cucumber growth in the natural soil system. (A) Representative images of the positive effect of increasing richness of MR consortia on cucumber growth. Increasing the richness of moderately related (MR) consortia (2+37+43+SQR9) stepwise improved the cucumber shoot height (B) and shoot dry weight (C) in the natural soil system. Different letters above the bars indicate significant differences (*p* < 0.01, *t test*, n=9, mean±standard deviation).

Fig. S5 Measurement of five PGP traits of *Bacillus* strains used for building HR and MR consortia in Fig. 5B and 5C. A, NH_3_ production; B, phosphate solubilization ability; C, IAA production; D, siderophore production; E, growth ability. We assigned each measurement a relative value between 0 and 1 based on the min-max measured values for each trait. The last “summary artificially” pentagons indicate the total PGP trait prediction for each *Bacillus* consortium.

Table S1: Topological properties of the gene cooccurrence networks and their respective identically sized random networks.

Table S2: Annotation information by using the NCBI database for each node in Figure 2E and 2F based on the sequencing fragment (490 bp) in gene co-occurrence network analysis. The similarity of the *gyrA* gene for each node to SQR9 was also present.

Table S3: Detailed information of swarming interaction phenotypes for *Bacillus* isolates from both SS and SS_SQR9 treatments.

Table S4: Detailed information of the components of highly related (HR) consortia and moderately related (MR) consortia.

