## Supplementary figures and images for "*Bacillus velezensis* SQR9 sways the rhizosphere community and its sociality toward cooperation and plant growth promotion"

### Supplemental Figure S1

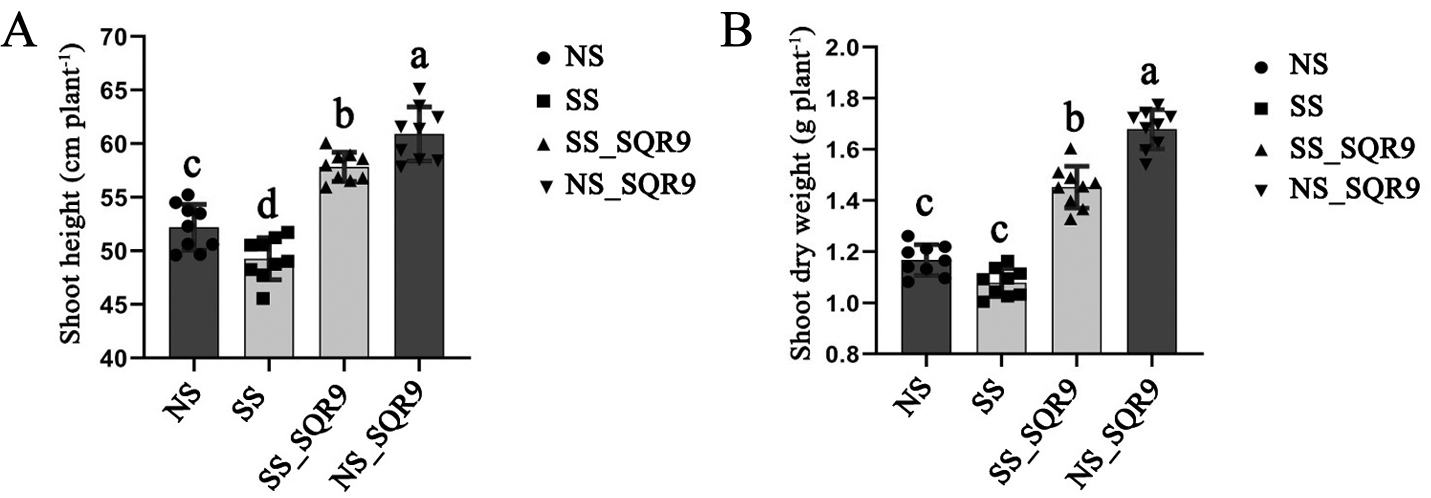

### Supplemental Figure S2

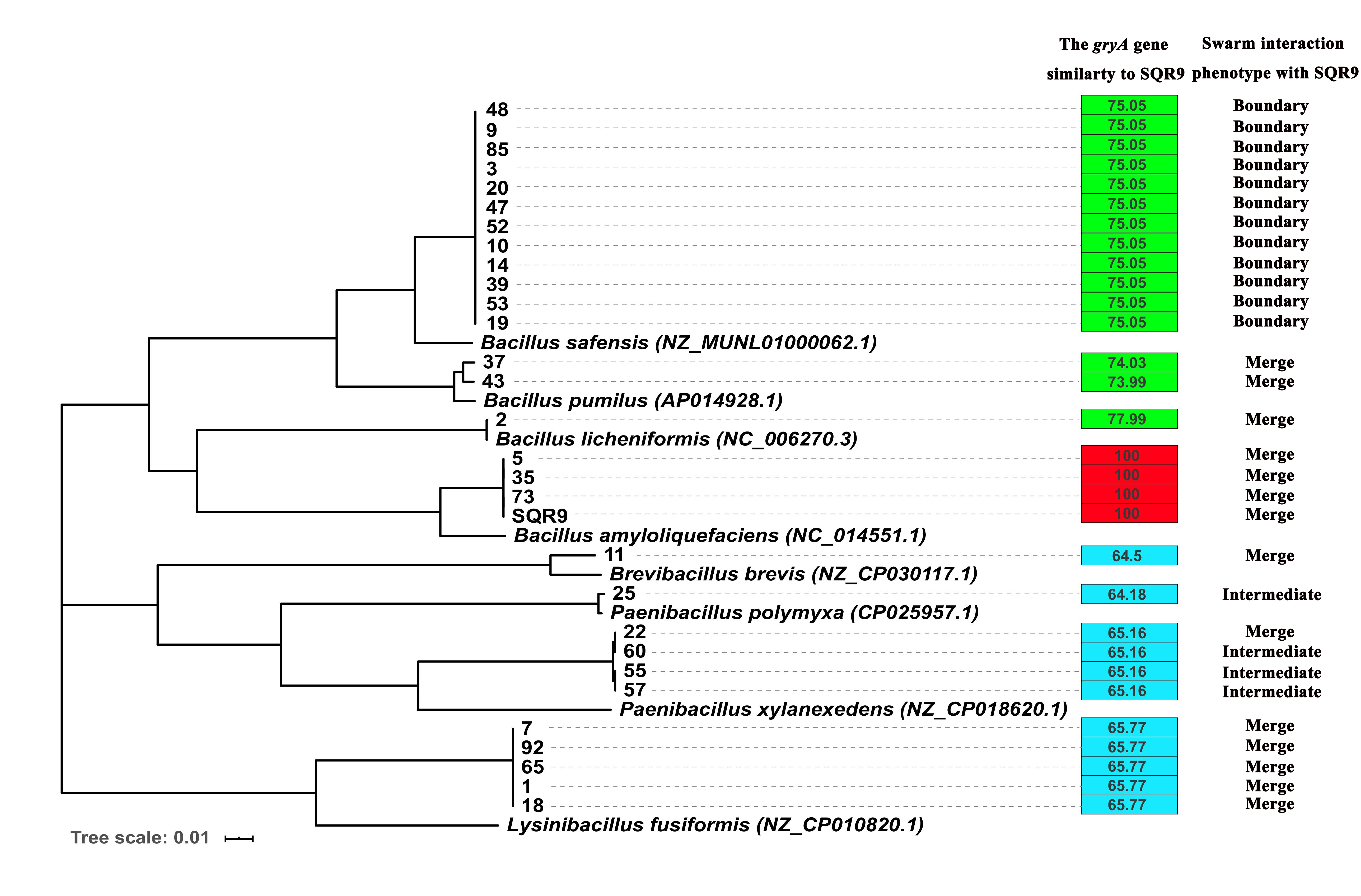

### Supplemental Figure S3

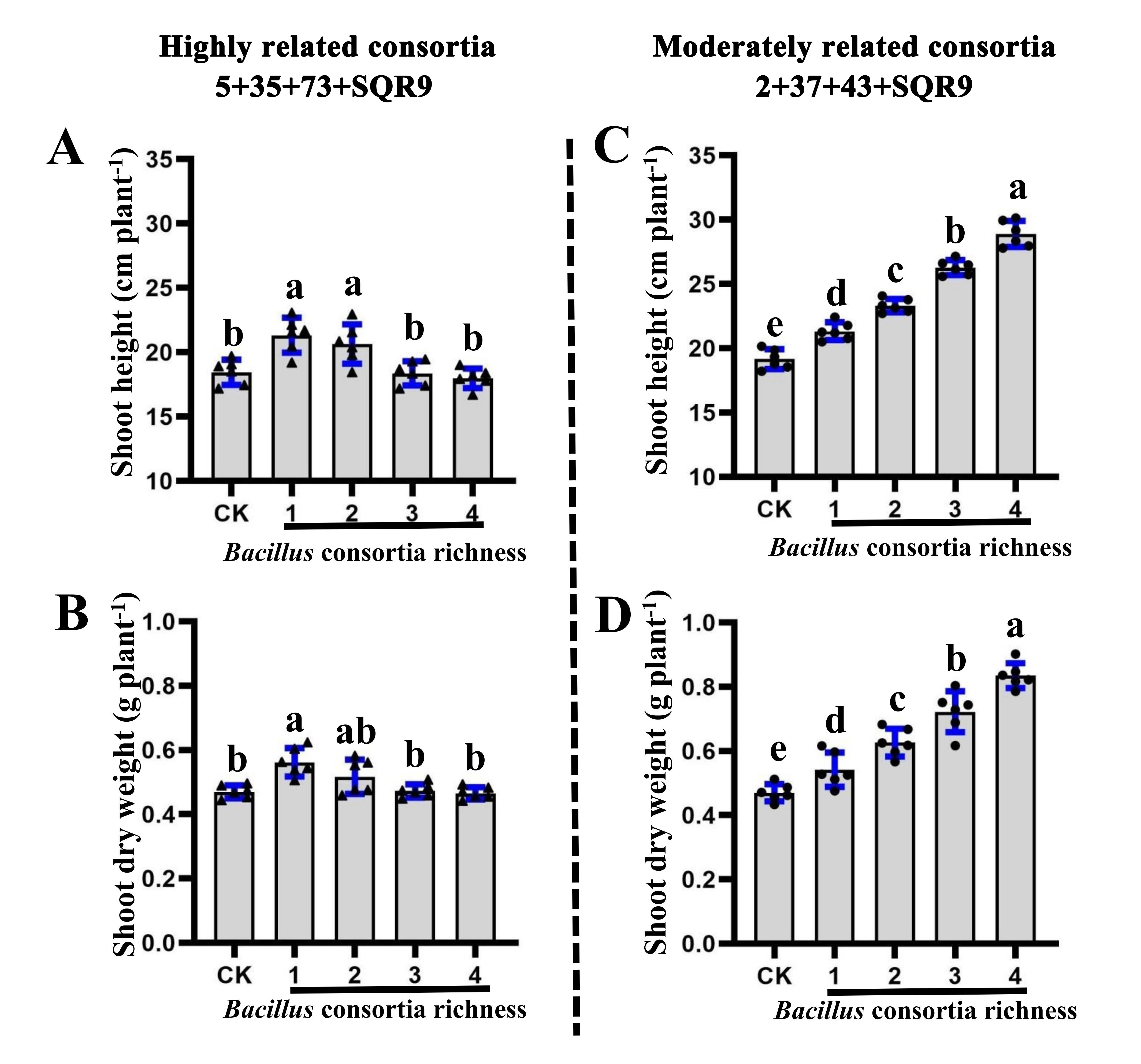

### Supplemental Figure S4

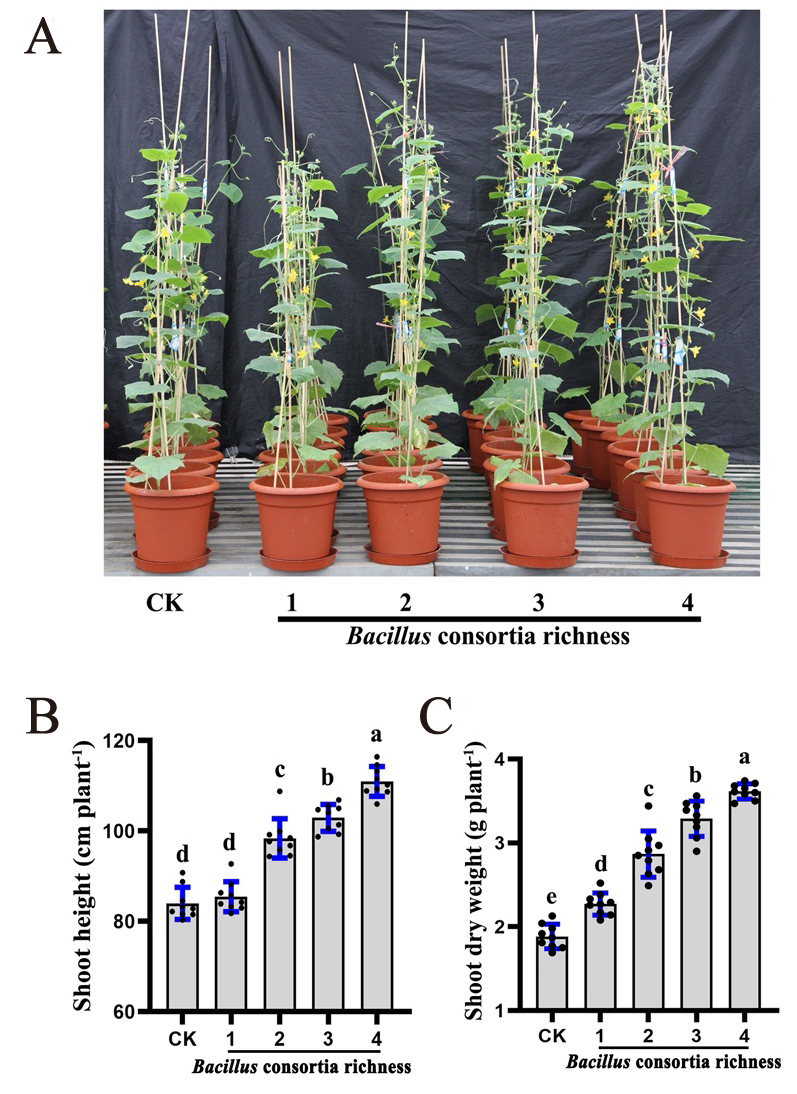

### Supplemental Figure S5

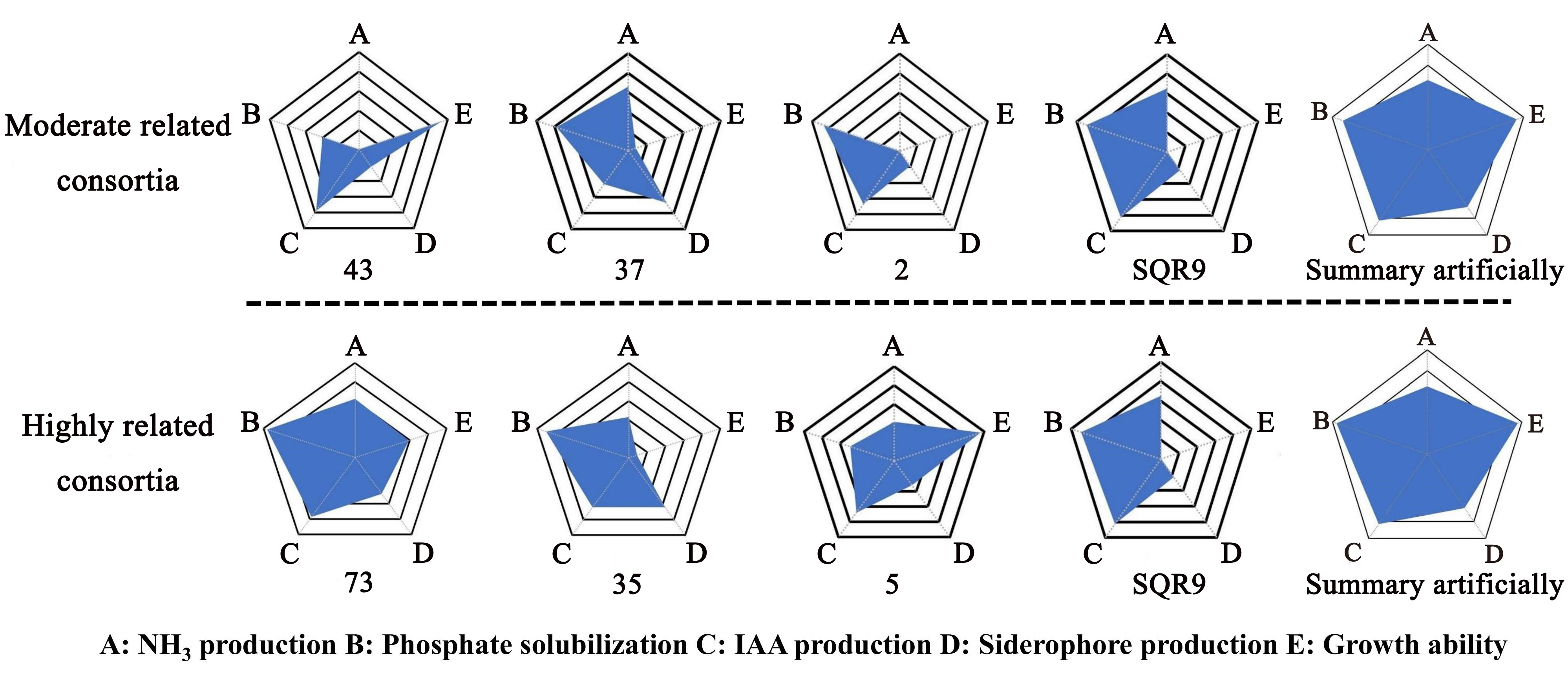
