## Supplemental Table S1 for "*Bacillus velezensis* SQR9 sways the rhizosphere community and its sociality toward cooperation and plant growth promotion"

Table S1. The topological properties of the gene co-occurrence networks and their respective identically sized random networks

| Network Indexes | non-SQR9-inoculated | SQR9-inoculated |
| --- | --- | --- |
| *Empirical Networks* |  |  |
| Total nodes | 453 | 294 |
| Total links | 5899 | 13103 |
| Average degree (avgK) | 26.04 | 89.14 |
| Average clustering coefficient (avgCC) | 0.689 | 0.836 |
| Average path distance (APD) | 3.855 | 2.226 |
| Modularity (M) | 1.116 | 1.175 |
| Graph density (GD) | 0.058 | 0.304 |
| *Random Networks* |  |  |
| Average clustering coefficient (avgCC) | 0.452±0.018 | 0.5178±0.037 |
| Average path distance (APD) | 1.241±0.012 | 1.025±0.024 |
| Modularity (M) | 0.342±0.005 | 0.514±0.013 |

Obviously, inoculation with strain SQR9 into soil exhibited a strong impact with a reduction in the total nodes (decreased from 453 to 294), however, inoculation with strain SQR9 significantly increased the total links (positive or negative correlations), average degree (avgK), average clustering coefficient (avgCC), graph density (GD), indicating that inoculation with strain SQR9 re-assembled rhizosphere soil microbiome and further improved the correlation degree among the bacteria community in rhizosphere (Table S1). In addition, inoculation with strain SQR9 into soil also exhibited a slight increase in modularity indices (Table S1). In summary, inoculation with strain SQR9 into soil increased the complexity of gene co-occurrence networks.
