## Supplemental results and methods for "*Bacillus velezensis* SQR9 sways the rhizosphere community and its sociality toward cooperation and plant growth promotion"

College of Resources and Environmental Sciences, Nanjing Agricultural University, Nanjing, 210095, P.R. China; Department of Microbiology, Biotechnical Faculty, University of Ljubljana, Ljubljana, Slovenia

**Supplemental results**

**Supplemental Results 1**

**The non-metric multidimensional scaling (NMDS) analysis of 16S rRNA and *gyrA* amplicon sequencing data**

To explore what extent the introduced strain SQR9 influences the composition of the rhizosphere microbiome, we used the nonmetric multidimensional scaling (NMDS) analysis of 16S rRNA and *gyrA* amplicon sequencing data. Results showed that NMDS of 16S rRNA failed to reveal the effect of SQR9 inoculation on bacterial communities (NS versus NS_SQR9) (Statistical ANOSIM R value = 0.241, *p =* 0.031, Fig 2C). In contrast, a significant shift (*p* < 0.01) in the composition of *gyrA* gene was observed in Fig. 2D (NS versus NS_SQR9). Moreover, the composition of the *gyrA* gene varied extensively between SQR9 treated or untreated soils (both natural and sterilized) as *gyrA* sequences of treated and untreated samples are clearly separated on the horizontal (Statistical ANOSIM R value = 0.861, *p <* 0.001, Fig. 2D), as well as on vertical axis (Statistical ANOSIM R value = 0.577, *p =* 0.018, Fig. 2D). Overall these results support the conclusion that SQR9 shifts the composition of indigenous bacterial community and that these shifts are more easily detected at the lower than at the higher taxonomic level, where the diversity is more broadly distributed.

**Supplemental Results 2**

**Network Analysis of *gyrA* co-occurrence patterns**

Since *gyrA* analysis revealed more prominent shifts in rhizosphere bacterial communities, we applied network analyses of the *gyrA* gene cooccurrence patterns to further address interactions between SQR9 and the rhizosphere community and its impact on community assembly. To determine cooccurrence patterns, we used the network analyses based on the samples with strong and significant correlations (spearman’s correlation coefficient R > 0.80, *p <* 0.01, two-sided tests). The network analyses were performed for untreated samples (NS and SS) and SQR9 treated (NS_SQR9 and SS_SQR9) samples separately. Results showed that a) inoculation of SQR9 exhibited a strong impact with a reduction in the total nodes but an increase in links (Fig. 2E, 2F and Table S1); b) the average degree, average clustering coefficient, modularity, and graph density indices were higher in the SQR9 inoculated network (Fig. 2E, 2F and Table S1); c) the modularity indices increased from 1.116 to 1.175 with SQR9 inoculation (Table S1). In addition, SQR9 enriched the proportion of *Bacillus* species in the rhizosphere from 6.4% to 49.83% (Fig. 2E and 2F). Overall, our results confirming that SQR9 affects the composition and co-occurrence pattern of rhizosphere bacteria community.

Moreover, color of the nodes further depicts the level of relatedness between SQR9 *gyrA* and other *gyrA* sequences. For example, nodes that were highly related to SQR9 (98 ≤ genetic distance < 100) are colored red (Fig 2E and 2F). In the untreated soil network, a large proportion (89.18%, 404 out of 453 nodes) of nodes were represented by phylogenetically distant members, observed as blue nodes (50 ≤ genetic distance < 60 to SQR9) (Fig. 2E and Tables S2). In the SQR9-treated sample (Fig. 2F), beside SQR9, which was the most abundant strain in the network (indicated by black arrow and letter A in Fig. 2F), two modules were significantly enriched: Module 1 with highly related members (98 ≤ genetic distance < 100, red nodes) representing genotypes of *B. velezensis* species (Table S2) and Module 2 with moderately related members (70 ≤ genetic distance < 80, green nodes) (Fig. 2F). In contrast, distant relationship members (50 ≤ genetic distance < 60, blue nodes) were drastically reduced (50.34%, 148 out of 294 nodes) in SQR9-inoculated sample (Fig. 2F and Tables S2). Overall, these results potentially indicate that SQR9 engages in interactions with members of the bacterial community in the rhizosphere and consequently decreases the abundance of some members and promotes the growth of others. Increased number of nod connections within clusters suggests that SQR9 enhanced interactions between genotypes within modules (Figure 2F, Table S1 and S2).

**Supplemental Results 3**

***B. velezensis* SQR9 increased compatibility of the *Bacillus* community in the rhizosphere**

In order to associate different swarming patterns from SS and SS_SQR9 rhizosphere to the relatedness of interacting strains inside each tratment we determined the phylogenetic distance between *Bacillus* strains from SS and SS_SQR9 treatments by reconstructing the minimum evolution trees based on their *gyrA* gene sequences. Results show that 30 *Bacillus* strains from the untreated SS rhizosphere show higher richness (15 species) (Fig. 4A) than SQR9 tretaed soil (8 species) (Fig. 4C). Moreover, the frequency of merging and boundary phenotype pattern changed after inoculation of SQR9. For *Bacillus* isolates in SS treatment, the boundary phenotype occurred between almost all strains within the arbitrary clusters (boundaries=125, merging=8) and between almost all strains within the cluster 1 (*subtilis*/*amyloliquefaens*/*mojavensis* group, boundary=102, merging=3), cluster 2 (*pumilus* /*sefensis* groups, boundary=6, merging=1), cluster 5 (*thuringensis* group, boundary=15, merging=0). Boundaries also dominated between closely related clusters 1, 2, 5 (boundary = 105). However, combinations with more distant species (cluster 3 and 4) often resulted in swarm merging (Fig. 4A and 4B). In contrast, among isolates from SS-SQR9 rhizosphere the merging phenotype dominated between strains within clusters (merging=92, boundary=28) and always among strains within the same species. Additionally, merging between clusters was also enriched (merging=164, boundary=52) (Fig. 4D). Overall, these observations agree with the hypothesis that SQR9 decreases the frequency of antagonists and promotes enrichment of compatible strains.

**Supplemental results 4**

**Social interactions and resource competition overpower *in vitro* determined plant growth promoting (PGP) traits for plant growth promotion**

To test whether plant growth promotion is affected by individual and cumulative plant growth potential of HR and MR strains, we further determined five PGP traits of each isolate (growth ability, production of IAA, siderophore, NH_3_ and phosphate solubilization) used in the consortia. Results showed that HR isolates’ individual PGP potential is higher that MR consortia isolates (depicted as blue surface area in the pentagrams) (Fig. S5), however, the artificial sum of PGP traits inside HR or MR isolates is similar (the blue area on the pentagram). Nevertheless, plant growth promotion was observed to be higher with the MR isolates (Fig. 5B and 5C) despite their lower individual potential (Fig. S5) to promote plant growth, indicating that ecological interactions, comprising sociality and resource competition play an important role in PGP.

**Supplemental Methods 1**

**Selection of strains for merging assays**

Rhizosphere soils from SS and SS_SQR9 treatments (see Figure 1A) were resuspended in 1 mL of sterile saline solution (0.9% NaCl), and the suspension was heated for 15 min at 80 °C to kill vegetative cells but preserve spores. Resultant spore suspensions were plated on tryptose blood agar base (Difco; Becton, Amresco, China, ID: 227300) and incubated for 24 h at 30 °C. Emergent colonies were streaked three times to obtain pure cultures, yielding 280 isolates. We selected 30 represent isolates for both SS and SS_SQR9 treatments, and the selection phenotypic criteria as follows: a, 16S rRNA nucleotide identity and three metabolic tests (the catalase test, the Voges-Proskauer test and anaerobic growth on agar)^1^ indicated it belongs to *Bacillus* and its related species; b, isolates can form biofilm in MSgg medium; 3, isolates can swarm on agar plate, ranked 30 top isolates according to its swarm ability were selceted from SS and SS_SQR9 treatments respectively.

**Supplemental Methods 2**

**Phenotypic characterization of *Bacillus* isolates from cucumber rhizosphere**

Five PGP phenotypic traits (NH_3_ production, growth ability, siderophore production, IAA production and phosphate solubilization ability) were tested for each individual *Bacillus* isolates which used to establish random communities (consortia) using rhizosphere isolates. In order to prepare the bacterial inoculant, one colony was picked and pre-grown overnight at 37 °C in LB, washed three times in 0.85% NaCl and adjusted to a density of 10^8^ cells mL^-1^.

Ammonia (NH_3_) production. Ammonia production was detected by using peptone water^2^. We grew all individual *Bacillus* isolates in 1.5% w/v peptone water for 2 days at 30 °C with shaking (150 rpm). Cell-free supernatants were added with 5% Nessler reagent. Nesslerization of sterile un-inoculated peptone water was served as reference. Colour change of supernatants from pale to deep yellow was determined at absorbance 425 nm. The amout of produced ammonia was measured throuhg a standard curve established by determining the different concentration (0-100 mM) of authentic amonia at absorbance 425 nm.

Growth ability. Growth of each *Bacillus* isolates were assessed in 200 μL Minimal medium-glucose-yeast (MGY) medium (glucose 5 g L^-1^, yeast extract 4 g L^-1^, NH_4_NO_3_ 1 g L^-1^, NaCl 0.5 g L^-1^, K_2_HPO_4_ 1.5 g L^-1^, KH_2_PO_4_ 0.5 g L^-1^, MgSO_4_ 0.2 g L^-1^, pH 7.0) in 96-wells microtiter plates. The initial inoculum size was set at OD_600_ value of 0.05. OD_600_ was measured every 30 minutes at 30 °C with Bioscreen C Automated Microbiology Growth Curve Analysis System (Growth Curve, USA). This assay was repeated three times.

Siderophore production. We grew all individual *Bacillus* isolates in MKB medium (K_2_HPO_4_ 2.5 g L^-1^, MgSO_4_ 7H_2_O 2.5 g L^-1^, glycerin 15 mL L^-1^, casamino acid 5 g L^-1^, pH 7.2) for 48 h at 30 °C with constant shaking (170 rpm). After centrifugation (10000 g for 5 min), the siderophore production was assayed using a modified method developed by Schwyn and Neilands^3^. Briefly, 0.5 mL of cell-free culture supernatant or deionized water as a control reference were mixed with 0.5 mL of CAS assay solution. After 2h of static incubation at room temperature, the OD_630_ of cell-free supernatants (A) and deionized water control (Ar) was then measured using a plate reader (SpectraMax M5) at room temperature. Siderophores induce a colour change in the CAS medium, which lowers the OD_630_ measurements, and the siderophore production can thus be quantified using the following formula: 1-A/Ar^4^. This assay was repeated three times.

IAA production. We grew all individual *Bacillus* isolates in liquid Landy medium (glucose 20 g L^-1^, L-glutamic acid 5 g L^-1^, KH_2_PO_4_ 1 g L^-1^, yeast extract 1 g L^-1^, MgSO_4_ 7H_2_O 0.5 g L^-1^, KCl 0.5 g L^-1^, MnSO_4_ H_2_O 5 mg L^-1^, CuSO_4_ 7H_2_O 0.16 mg L^-1^, FeSO_4_ 7H_2_O 0.15 mg L^-1^, L-phenylalanine 2 mg L^-1^, L-tryptophan 1 g L^-1^, pH 7.0) for 72 h at 22 °C in the dark with constant shaking (90 rpm). Bacterial cultures were then centrifuged (at 10000 g for 5 min) and IAA concentration of the supernatants (ng/mL) was measured with IAA ELISA Kit (R&D, Shanghai, China) following the manufacturer’s protocol^5-6^. This assay was repeated three times.

Phosphate solubilization ability. We grew all individual *Bacillus* isolates in NBRIP medium (glucose 10 g L^-1^, Ca_3_(PO_4_)_2_ 5 g L^-1^, MgCl_2_ 6H_2_O 5 g L^-1^, MgSO_4_ 7H_2_O 0.25 g L^-1^, KCl 0.2 g L^-1^, (NH_4_)_2_SO_4_ 0.1 g L^-1^, pH 7.0) for 7 days at 30 °C with constant shaking (170 rpm). After centrifugation (at 10000 g for 10 min) the soluble phosphate concentration (μg/mL) of the supernatant was measured using the molybdenum antimony colorimetric method. Measurements were replicated three times^7^.

Multifunctionality analysis. To predict how multifunction traits can be analytically related to the potential of soil isolates/communities to promote plant growth we assigned to each isolate a value for each trait. After measuring five tested ecological traits for each isolate we assigned each measurement a relative value between 0 and 1 based on the min-max measured values for each trait. The results are presented as the radar chart (pentagram) for each individual *Bacillus* isolate and communities. The sum of these ecological trait’s values was then used to speculate on overall multifuncionality of each isolate.
